# A Burst-Dependent Algorithm for Neuromorphic On-Chip Learning of Spiking Neural Networks

**DOI:** 10.1101/2024.07.19.604308

**Authors:** Michael Stuck, Xingyun Wang, Richard Naud

## Abstract

The field of neuromorphic engineering addresses the high energy demands of neural networks through brain-inspired hardware for efficient neural network computing. For on-chip learning with spiking neural networks, neuromorphic hardware requires a local learning algorithm able to solve complex tasks. Approaches based on burst-dependent plasticity have been proposed to address this requirement, but their ability to learn complex tasks has remained unproven. Specifically, previous burst-dependent learning was demonstrated on a spiking version of the XOR problem using a network of thousands of neurons. Here, we extend burst-dependent learning, termed ‘Burstprop’, to address more complex tasks with hundreds of neurons. We evaluate Burstprop on a rate-encoded spiking version of the MNIST dataset, achieving low test classification errors, comparable to those obtained using backpropagation through time on the same architecture. Going further, we develop another burst-dependent algorithm based on the communication of two types of error-encoding events for the communication of positive and negative errors. We find that this new algorithm performs better on the image classification benchmark. We also tested our algorithms under various types of feedback connectivity, establishing that the capabilities of fixed random feedback connectivity is preserved in spiking neural networks. Lastly, we tested the robustness of the algorithm to weight discretization. Together, these results suggest that spiking Burstprop can scale to more complex learning tasks and can thus be considered for self-supervised algorithms while maintaining efficiency, potentially providing a viable method for learning with neuromorphic hardware.

## 1. Introduction

The increasing energy demands of neural network-based artificial intelligence (AI) systems have created significant interest in developing more energy-efficient methods [1, 2]. Neuromorphic computing, which takes inspiration from the computational principles of the human brain, has emerged as a promising solution to reduce energy costs associated with AI systems. Two biological aspects, in particular, are thought to yield significant energy savings.

One is the event-based mode of communication [3, 4, 5]. Unlike traditional neural networks that rely on high-precision, continuous activations, SNNs operate using single-bit, binary activations, known as spikes. Running SNNs with low-precision parameters and high spatial and temporal sparsity levels contributes to this energy efficiency [1].

The other advantageous biological property is the co-localization of computing and memory which circumvents the von Neumann bottleneck [6, 7]. Current methods for training SNNs predominantly rely on conventional von Neumann computer architectures, with the learned synaptic weights transferred to spiking neuromorphic hardware to enable energy-efficient inference. While using spiking neuromorphic hardware for inference has demonstrated energy savings in certain applications [8], further energy efficiency can be achieved by training SNNs directly on hardware. Direct hardware training reduces energy consumption and allows for continual learning, essential for systems that need to dynamically adapt to changing environments. Achieving this requires new training algorithms that are compatible with the unique constraints of neuromorphic hardware.

The constraints presented by neuromorphic hardware for learning have many similarities with those present in biological systems. As such, a learning algorithm for spiking neuromorphic hardware must include certain characteristics which are thought to be important for biologically plausible learning. The most essential of these biologically plausible features of learning is local, event-driven synaptic plasticity [9]. This means that synaptic weight changes must be triggered by either presynaptic, postsynaptic or global events and can depend only on the states of the presynaptic neuron, postsynaptic neurons, a global state and the synapse itself. A local alternative to backpropagation is necessary to do on-chip learning of SNNs in neuromorphic hardware.

One promising group of local learning methods for training SNNs are burst-dependent, ‘Burstprop’ or BurstCCN learning methods [10, 11]. These algorithms are grounded in the widely observed burst-dependence of synaptic plasticity [12, 13, 14, 15] and synaptic dynamics which confers synapses with the ability to communicate spike-timing patterns differently to different targets. In hierarchical networks, Burstprop can approximate the backpropagation of error algorithm with single-phase local learning. It accomplishes this by an ensemble multiplexing [16] of error signals, which are transmitted backwards through the network and integrated by dendritic compartments to steer plasticity while the network processes feedforward information through somatic neuron compartments. Thus far, however, Burstprop methods have been limited to solving the exclusive or (XOR) problem in SNNs [10, 11]. For Burstprop to be considered viable for neuromorphic learning applications, it must be capable of learning more difficult tasks. Similarly, we currently don’t know if feedback alignment, a technique used to perform credit assignment without an artificial tying of the connectivity, preserves its function in SNNs using local and online plasticity rules.

In this work, we improve and test current Burstprop methods, allowing it to scale to more difficult learning problems. We consider two improvements: 1) a regularizing feedback connection from the soma to the dendrite of every neuron and 2) communication of two types of bursts, allowing for the multiplexing of signed errors. We evaluate our algorithm using the MNIST digit classification and other related benchmarks (EMNIST and FMNIST) and show that it can successfully learn this task. We also investigate the impact of the choice of feedback connectivity and weight discretization. The results support that our algorithm is a viable method for supervised learning of SNNs for neuromorphic implementation, a first step for implementation of more elaborate algorithms such as large unsupervised, self-supervised or deep-reinforcement-learning models.

## 2. Background

### 2.1. Learning in SNNs

Multiple methods for training SNNs have been developed. We first distinguish methods relying on non-local plasticity rules from those using local ones. The local plasticity rules can be further separated into burst-dependent plasticity and spike-timing-dependent plasticity.

#### 2.1.1. Non-Local Learning Methods

Nonlocal learning methods for SNNs include ANN-SNN conversion [17, 18, 19, 20] and backpropagation-through-time (BPTT) [21, 22]. With ANN-SNN conversion, a rate-based neural network is trained using the backpropagation algorithm which is then converted to a spiking neural network. In real applications, imperfect conversions will strongly alter performance, but this can be remedied with further fine-tuning [8]. With BPTT, backpropagation is performed on an SNN by using surrogate gradients to propagate the gradient through the spiking nonlinearity. To backpropagate gradients through time, the gradient must keep track of the network state at multiple time steps of past activity for each time step. This algorithm is therefore highly nonlocal (but local-only approximations have been derived [23]). These methods do not address the problem faced with training neuromorphic online, we will use these methods to provide a point of reference for the performance metrics.

#### 2.1.2. STDP Learning Methods

Spike-time-dependent plasticity (STDP) alters connectivity according to the relative timing of spikes in the presynaptic and postsynaptic neurons and is therefore fully local. One local learning algorithm, proposed by Kappel et al. [24] combines a STDP plasticity rule and winner-take-all dynamics and recurrent excitation to approximate hidden Markov model learning in SNNs. Relatedly, Diehl and Cook [25] combined STDP with winner-take-all dynamics to train a non-recurrent network. Reading out the resulting representation with a supervised plasticity rule, they showed that a single layer could achieve 95% accuracy on MNIST. As only the readout is supervised, this method is sometimes referred to as unsupervised. Another STDP approach combines a trace set by relative coincidence and a global reinforcement by reward-associated signals [26, 27, 28], but such three-factor learning rules tend to scale poorly on problems having a high number of dimensions. Also, how these STDP approaches can scale to multiple layers in a hierarchy remains unknown.

#### 2.1.3. Burstprop Learning Methods

Another class of local learning algorithms depend on separate somatic and dendritic compartments to represent both feedback error signals and inference signals within the neuron, often through a rate-based representation [29, 30, 31, 32]. For spiking neural networks, burst-dependent algorithms exploit the target-dependence of short-term plasticity [33, 34, 35], dendrite-dependent bursting [36, 37] and burst-dependent plasticity [13, 12, 14, 15] to perform credit assignment in a way that approximates backpropagation [10, 11, 38]. These burst-dependent learning methods, referred to as ‘Burstprop’ algorithms, can in principle allow for multilayer local learning in SNNs.

The original formulation [10] of the algorithm learns online but with signals coming in two phases. An initial phase in which synaptic plasticity was disabled was required to determine each neuron’s baseline burst probability which was then used in the second phase to calculate synaptic plasticity, violating temporal locality. A second formulation of the algorithm (called BurstCCN [11]) adapted the learning rule for single-phase learning by including a second feedback pathway that communicates spike signals such that a baseline burst probability can be preset, avoiding the need for two phases. Addressing complex problems with these algorithms has been almost entirely limited to rate-based networks. Spiking proof-of-principles were done on the XOR problem.

### 2.2 Weight-Transport-Free Learning

Approaches approximating the backpropagation of error algorithm face a problem known as the weight-transport problem. To accurately approximate the gradient, the feedback pathways must have synaptic weights that are symmetric to feedforward weights (symmetric feedback). Such direct weight-transport involves nonlocal communication weight values between synapses and must be avoided for learning in spiking neuromorphic hardware. Many solutions have been proposed [39, 40, 41, 42, 43, 10, 44, 45], including imposing random but static feedback connectivity (feedback alignment [39, 40]) and learning rules that seek to transport the weights by exploiting correlations in the activity patterns [42, 10, 44].

#### 2.2.1. Feedback Alignment Methods

It has been shown that feedback weights need not be initialized as symmetrical to feedforward weights, nor to be kept tightly symmetrical throughout learning to effectively train networks using backpropagation [39]. As long as feedforward weights are initialized within 90° of feedback weights, feedforward weights trained with backpropagation tend to align with feedback weights, leading to accurate credit assignment and effective learning. Since feedforward synapses must align with feedback synapses, this method is referred to as feedback alignment (FA). Building upon this work, direct feedback alignment (DFA) uses random feedback weights directly connecting network outputs to each layer and has also been shown to be effective for learning with backpropagation [40]. While these methods are effective for training shallow, rate-based neural networks, their efficacy tends to decrease with increasing network depth [46, 47, 48].

The question of whether these techniques also scale to spiking neural networks has been partially addressed in Samadi et al. [41], where a spiking feedforward inference phase was combined with an analog learning phase that follows the backpropagation of error algorithm. Zenke and Ganguli [49] have similarly paired a spiking inference phased with analog, non-local, BPTT learning. Relatedly, Zhao et al. [50] have studied SNNs receiving direct but analog gradient information. Xing et al. [51] have used a signed spiking communication for the feedforward inference phase, but again paired with analog BPTT communication of gradients. In Payeur et al. (2021) [10] a fully spike-based communication of both feedforward and learning signals, but the feedback alignment approach was only applied to the XOR problem. The present work focuses on MNIST in a fully spiking communication of both feedforward and learning signals.

#### 2.2.2. Learned Feedback

Several methods implement a learning rule on feedback synapses, allowing them to align with feedforward weights. With the Kolen-Pollack method [52], weight symmetry is achieved by assigning the same plasticity rule to feedforward and feedback synapses and including a weight decay term. While this is effective in aligning weights, the necessity of weight decay may be a problem for some learning methods. Phaseless Alignment Learning (PAL) aligns feedback weights to feedforward weights by learning feedback weights from neuronal noise [44] outside of the presentation of examples. Another algorithm proposed by Burbank [53] uses temporally opposed STDP learning rules on feedforward and feedback synapses to successfully train a spiking autoencoder network. Similarly, Payeur et al. [10] provide a spike-based learning rule for learning the alignment of feedback synapses, which have not been tested on SNNs. These methods form a conceptual rationale for using implementing symmetry as a partially accurate bypass of the learned feedback method.

## 3. Methods

### 3.1. Neuron and Synapse Model

#### 3.1.1. Burst Coding

We consider a neural code in which neurons are thought to emit spikes in two different modes: either bursts or single spikes. Bursts are a set of action potentials emitted at a high frequency, typically 100 Hz [15]. There are at least 2 spikes per burst, but the number of spike per burst is not a requirement for its class. Single spikes are relatively isolated action potentials. Both bursts and single spikes are considered events. We note that it in such a code, it has been shown that a small number of spikes per bursts and a low burst probability is maximizing information transmission [54].

In what follows, we will also be introducing the notion of two different types of bursts. We consider that a simple interpretation of such a code is to consider long bursts (e.g. bursts having more than 5 or 6 spikes) and short bursts (e.g. bursts having 4 spikes or less). While there is only limited evidence that different bursting modes (long and short) are regulated in the brain [15], we are not aware of any evidence to support the fact that such modes of firing would be regulated by different types of inputs to the same dendrite, we consider it is mainly an interesting avenue for neuromorphic hardware implementations. Because of their relationship with the input signal, we will label these different types of bursts ‘positive’ and ‘negative’ bursts.

It is assumed based on experimental observation [54, 10] that inputs to the soma are driving the generation of any event, wether single spike or burst. The net inputs to the dendrite regulates whether this event is a burst or a spike in a probabilistic way, such that a population of neurons can multiplex two independent streams of information in its time-dependent event rate and time-dependent burst fraction. Evidence of this multiplexing has been observed experimentally [55].

#### 3.1.2. Somatic Compartment

Each neuron’s soma follows Leaky Integrate and Fire (LIF) dynamics [56]:

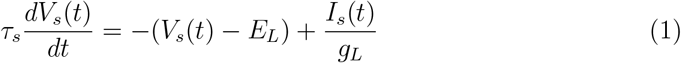

where *V*_*s*_(*t*) represents the somatic membrane potential, *τ*_*s*_ the membrane time constant, *E*_*L*_ the resting potential, *g*_*L*_ the leak conductance, and *I*_*s*_(*t*) the input current. LIF models have been shown to be quantitatively accurate models of many different cell types especially when integrating spike-frequency adaptation and nonlinear dendritic compartments [57, 58, 59, 60, 61]. The two-compartment LIF model without adaptation was chosen so as to implement the simplest set of dynamics able to capture credit assignment by burst-dependent plasticity. In our experiments, the somatic membrane time constant was set to *τ*_*s*_ = 10 ms. The neuron fires an event when *V*_*s*_(*t*) exceeds the threshold potential *V*_threshold_, following which *V*_*s*_(*t*) resets to *V*_reset_:

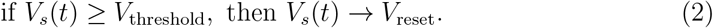

Rescaling to dimensionless variables, we define *Ĩ*_*s*_ = *I*_*s*_*/g*_*L*_(*V*_threshold_ − *E*_*L*_) and 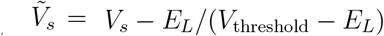, equation (1) becomes

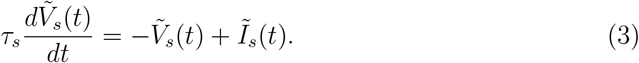

Every time the somatic voltage crosses the threshold, an event is emitted. As mentioned in the previous section, this event can be either a single spike or a burst, as determined by the dendritic state (see Sect. 3.1.4).

#### 3.1.3. Eligibility Trace

We use eligibility traces, taken as the exponential moving average of the event train *Ē*_*j*_ (*t*), to indicate the eligibility of individual synapses connecting from a lower-order neuron *j* to some other higher-order neuron:

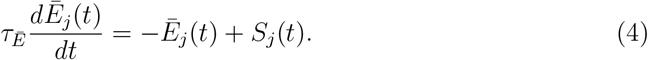

Here the event train 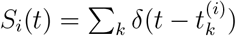, where 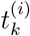 is the *k*^*th*^ event time from neuron *i*. The eligibility trace time constant was to 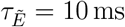.

#### 3.1.4. Dendritic Compartment and Burst Generation

Dendrites come with different properties to the neuron’s soma [62]. Here the dendritic compartment refers to an apical-like dendrite with a segregated membrane potential that can nonlinearly integrate feedback information to modulate neuron burst generation [36, 63, 60, 64]. It follows leaky integrator dynamics, using dimensionless inputs and membrane potential:

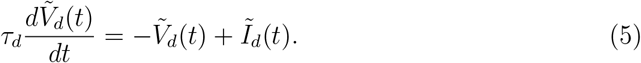

where 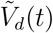 represents the re-scaled dendritic membrane potential, *τ*_*d*_ is the dendritic membrane time constant, and *Ĩ*_*d*_ is the re-scaled net dendritic input. We set the dendritic membrane time constant to *τ*_*d*_ = 10 ms.

There is no threshold in the dendritic compartment, instead, every time the soma emits an event, the state of the dendrite is used to determine if this event is a burst or a single spike. We consider two bursting models.

In the unsigned bursting model, the probability that an event is made into a burst is proportional to the dendritic potential 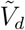 [65], we use a sigmoidal relationship 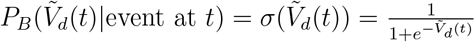.

In the signed bursting model, two types of bursts can be emitted (e.g. long bursts and short bursts, see Burst Coding Section). Either type of burst can only be generated upon the firing of an event from the soma. Given a somatic event, a burst is generated according to the symmetric function 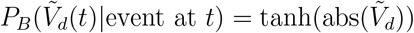. If a burst is said to be generated, the sign of *V*_*d*_(*t*) is used to determine its type. If 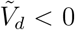, a short burst is generated. If on the other hand 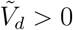, a long burst is generated. Because they represent the sign of the dendritic input, we refer to short and long bursts as negative and positive bursts, respectively. In this model, burst generation triggers a reset of the dendritic potential to the baseline, comparable to the somatic membrane potential reset upon a neuron spiking. This adds some refractoriness allowing the algorithm to better approximate backpropagation.

### 3.2. Network Architecture

The SNNs consist of an input layer of 784 units, corresponding to the number of pixels in each sample image in the MNIST-like datasets. There are 1, 2, 3 or 4 hidden layers consisting of 100, 200, 400 or 800 hidden neurons as specified in the figures and associated text. The output layer of 10 neurons corresponds to the 10 classes of the MNIST dataset. Except for input units, each unit follows the two-compartment model described above. The network inputs (see Sect. 3.8), as well as any connection between hidden layers or from the hidden layer to the output layer follow the fully connected configuration. Weights are allowed to be positive or negative. The label corresponding to the network’s class is used in calculating the network’s output error and teaching signal (see Sect. 3.4.1).

### 3.3. Synaptic Communication and Multiplexing

In our fully connected hierarchical SNNs, the credit-carrying information is communicated by multiplexing feedforward signals using events and feedback error signals using bursts. For the feedforward signals, we assume that synapses operate instantaneous changes in postsynaptic membrane potential upon every event. The input 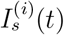 to the soma of neuron *i* is defined by the equation:

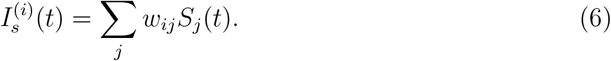

In this equation, *w*_*ij*_ is the feedforward synaptic weight from lower-order neuron *j* to higher-order neuron *i*.

#### 3.3.1. Unsigned burst Communication

For the feedback signals, we again assume that synapses operate instantaneous changes in postsynaptic membrane potential, but upon every burst. We further enforce that these feedback connections target the apical dendrite and therefore only affect the dendritic input 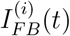 of a given neuron *i*. In the unsigned burst model, we have

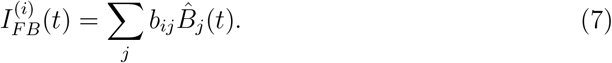

where *b*_*ij*_ are the feedback synaptic weight from higher-order neuron *j* to lower-order neuron *i* and 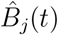 is the burst train of neuron *j*. Other, locally generated currents will combine with this feedback signal, see Sect. 3.4.2.

#### 3.3.2. Signed burst Communication

In the signed burst model, we assume the type of burst can dictate the sign of the post-synaptic effect:

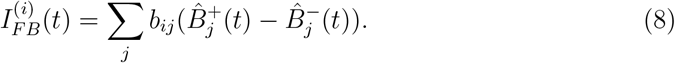

We note that since the same feedback weight *b*_*ij*_ operates on two distinct outputs of a neuron, this model requires weight transport.

#### 3.3.3. Multiplexing

In both signed and unsigned bursting models, when a neuron fires an event, the probability of that neuron sending a positive or negative error signal backward in the network is modulated by the dendritic potential of the neuron, *V*_*d*_, as demonstrated in Figure 2. Similarly, the event rate is proportional to the integrated somatic input (proportional to *V*_*s*_ if there were no reset). Thus the burst probability and the event rate represent two different signals simultaneously. Since each neuron integrates error signals from all higher-order neurons in its dendritic compartment and through the burst generation mechanism, the burst probability signal represents the signal that started with the input onto the apical dendrites at the top of the hierarchy and then backpropagates through it. This allows for the simultaneous online communication of credit-carrying error information from higher to lower-order neurons and feedforward, inference signals from lower to higher-order neurons.

### 3.4. Teaching Signals and Target Rates

For dendritic inputs to be conceived as providing error signals, we inject a teaching signal at the top of the hierarchy (Sect. 3.4.1) and a unit-specific regularisation signal (Sect. 3.4.2), which come together in some layers but not others (Sect. 3.4.3).

#### 3.4.1. Teaching Signals

Teaching signals are used for supervision, applying them to the dendritic compartments of output neurons. These signals are derived from the exponential moving average of the neuron’s firing rate, *Ē*(*t*), using the same time constant as for the eligibility trace (see Sect. 3.1.3). The teaching signal received by the dendrites at the top of the hierarchy, *T*_*i*_(*t*) is calculated as the derivative of the local squared error, *ε*_*T*_ (*Ē*_*i*_(*t*), *Ê*_*i*_), of the neuron’s instantaneous firing rate, *Ē*_*i*_(*t*) with respect to the neuron’s target rate, *Ê*_*i*_. For the neuron corresponding to the correct output class, a high target rate of *Ê*_*i*_ = 200 Hz is set, and for all other neurons, a low rate of *Ê*_*i*_ = 20 Hz is used. With this target, the teaching signal becomes

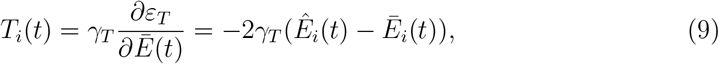

where *ϵ*_*T*_ corresponds to a square error loss. Here *γ*_*T*_ controls the strength of the teaching signal. This differential in target rates will backpropagate and control synaptic plasticity through the network, ensuring the correct output neuron becomes more active while other output neurons are suppressed. We found that a teaching signal strength of *γ*_*T*_ = 10^−3^ produced a strong enough teaching signal to steer plasticity while avoiding error signal saturation. Unlike in previous implementation of burstprop [10] we did not force the teaching signal to come after a delay, but the 100 ms was long enough to ensure that inference signal had ample time to reach the output layer and the associated gradient had ample time to reach the first hidden layer.

#### 3.4.2. Hidden target rates

Similar to the teaching signals applied to output neurons, hidden neurons receive a regularization signal that we model as a self-connection, such that 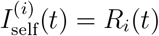 for all neurons except input neurons and output neurons. We use a uniformly low target rates *Ê* = 20 Hz:

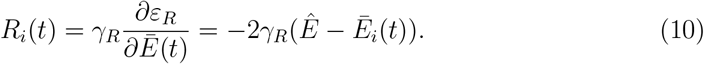

for *ϵ*_*R*_ a squared error loss. This *R*_*i*_(*t*) affects the apical dendrite of the same layer as the smoothed spike train *Ē*_*i*_(*t*), hence the self connection. We set *γ*_*R*_ = 10^−5^ ≪ *γ*_*T*_ such that the regularization does not suppress the error information due to the teaching signal. This approach regulates the firing rates of hidden neurons, thereby promoting sparsity and stability in the network’s learning process.

#### 3.4.3. Dendritic Inputs

Putting this together, we have that output units receive an dendritic input equal to the teaching signal and the regularizing signal

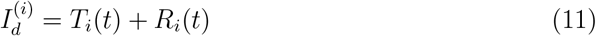

where *i* is the label of one of the output units. The hidden units receive a combination of the burst-coded feedback signals and the regularization signal:

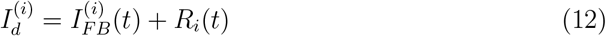

where *i* is the label of one of the hidden units. Input units are modeled as having no dendrites.

### 3.5. Plasticity Rule

Our algorithm utilizes a local spike-based learning rule to approximate backpropagation-based gradient descent. The synaptic weight changes are calculated as

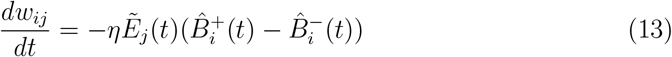

for the signed bursting model, and

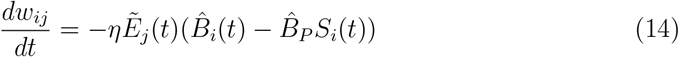

for the unsigned bursting model. Here, *B*_*P*_ is the baseline burst probability. The parameter *η* represents the learning rate and was set to *η* = 10^−4^ to achieve stable learning. We note that in the symmetric feedback scenario, the update of the feedforward weights *w*_*ij*_ also triggers an update in the feedback weight onto dendrites *b*_*ji*_.

For the signed bursting model, synaptic weight changes, occur upon postsynaptic burst generation, with the sign of the weight change determined by the type of burst. For the unsigned bursting model negative weight changes occur upon postsynaptic spikes and positive weight changes occur upon postsynaptic bursts. In both models, the magnitude of plasticity is proportional to the presynaptic neuron’s eligibility trace and the postsynaptic event rate. Plasticity, then, depends on three factors: the activity of the presynaptic neuron (via the eligibility trace), the postsynaptic neuron’s activity (triggering plasticity), and the integrated upstream error (via the burst fraction or burst type fraction). These three factors are essential to emulate the backpropagation of error algorithm, where weight adjustments of a given unit depend on three factors: the incoming activity, the derivative of the unit’s activity with respect to the sum of its inputs, and the integrated downstream error.

It is worth noting the distinct behaviour of signed and unsigned conditions in the absence of any postsynaptic error 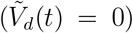. For the unsigned burst model, this condition results in a burst probability of 0.5. Because 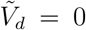 corresponds to the absence of dendritic input, we call the resulting output burst probability the baseline burst probability. In the associated plasticity rule, the parameter, *B*_*P*_ = 0.5, is chosen to match this baseline burst probability, such that there is a balance between positive and negative weight changes. We note that although this ensures that Δ*w* = 0 on average when 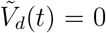, the fluctuations around this can cause drifts in the learned state over time. For the signed burst model, plasticity is more stable because the probability of burst generation is 0 when 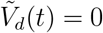 so no plasticity can occur.

### 3.6. Low-Resolution Weights

To implement low-resolution synaptic weights, *N*^2^ − 1 evenly spaced possible weight values were determined ranging from -1 and 1, where *N* is the weight resolution in bits. The magnitude of the interval between possible weight values is referred to as *ϵ*. This low resolution was imposed on initial weights and required an adaptation of the plasticity rule.

Initial weights (see Sect. 3.7) were rounded to the nearest possible weight value. During learning, an additional plasticity accumulator variable is added for each synapse.

Plasticity is calculated according to the learning rule and added to the accumulator variable. Once a synaptic accumulator reaches *±*0.5*ϵ*, the synaptic weight potentiates or depresses by *ϵ* and the accumulator variable weight resets. In this way, the network can learn with low-resolution weights while still having an effective learning rate of less than *ϵ*.

### 3.7. Weight Initialization

Feedforward synaptic weights were initialized from a normal distribution with standard deviation proportional to the square root of the number of presynaptic neurons to ensure stable learning [66]. It was found that to ensure error signals do not saturate, feedback weights had to be scaled by the number of higher-order neurons divided by the number of lower-order neurons in the layers connected by these weights. For the symmetric feedback implementation, feedback weights are aligned to feedforward weights. For FA and DFA, feedback weights were initialized from a normal distribution with the standard deviation scaled according to the number of higher-order neurons divided by the number of lower-order neurons while ensuring initial feedback weight matrices are within 90° of feedforward weights, as is done in [39].

### 3.8. Input Encoding

In our study, we encoded the MNIST Digit Classification Dataset images for a spiking neural network. The dataset consists of 60,000 training and 10,000 test grayscale images, each 28×28 pixels [67]. The images were normalized so that the average pixel intensity of each image was constant. Pixel intensities, ranging from 0 to 1, were linearly mapped to firing rates between 20 Hz and 200 Hz following a Poisson process. Accordingly, each pixel in each image was converted into an input unit spike train of a duration of 100 ms.

### 3.9. Simulation Methods

We simulated the SNNs using a CPU with Python. Neuron dynamics were simulated using the forward Euler method with a 1 ms time step. For each time step, the network was updated from the input layer to the output layer. Feedforward inputs to a layer were taken from the lower-order outputs at the same time step and feedback inputs to a layer were taken from the higher-order outputs at the previous time step.

#### 3.9.1. Training

During training, samples from the MNIST dataset are presented to the network sequentially. Teaching signals are applied continuously, and plasticity is engaged throughout the 100 ms. Neuronal state variables 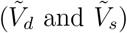, eligibility traces 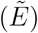 and rate estimates (*Ē*) are reset to zero at the end of each sample presentation.

#### 3.9.2. Testing

During testing, test samples are presented the same way as in the training phase, but there is no inputs to the dendrites and plasticity is off. The most active output neuron over the sample duration decides the network’s predicted class.

#### 3.9.3. Test in different conditions of initial sparsity

Different sparsity of connections were studied by initializing non-zero weights with different probability from 0.0 (no sparsity) to 0.8 (high sparsity). More specifically, we used the dropout function in Pytorch to set initialized weights to be zero with probability *p* layer by layer both for forward weights and backward weights. We note that the probability *p* does not correspond to the sparsity of the trained network as connections were allowed to change their weights from zero activity. Simulations where sparsity was changed were different set of simulation carried not only on the MNIST dataset but also on EMNIST’ and FashionMNIST. In these simulations, the number of training samples and test samples are 10000 and 1000.

### 3.10 Backpropagation-Through-Time

We compare our results with a backpropagation-through-time method using surrogate gradients [21]. We use the sigmoid function to approximate the spike discontinuity in the backward pass. The neurons follow the same dynamics, with the same parameters as the somatic neuron compartments previously described. We trained the network on the same input encoding, with each sample presented also for 100 ms and used a learning rate of *η* = 2 *×* 10^−4^ to balance stability and speed of learning.

## 4 Results

We trained SNNs with Burstprop [10] on a rate-coded version of the MNIST dataset. The idea is to use burst multiplexing to represent simultaneously the inference signal and the gradient signals, allowing for online learning with local learning rules. Burst multiplexing considers the neural code as being made with two types of events: either single spikes or bursts of high frequency action potentials. In a common implementation of this neural code, inputs to the dendrites controls the fraction of events that are busts while the inputs to the soma control the rate of event emission. The present implementation uses two-compartment neuron models with LIF soma and dendrite-dependent bursting. The bursts are not simulated as high-frequency spikes, but as labels to a spike train, that is, a marked point process. We will use the dendrite-encoded burst fraction to represent the gradient and the soma-encoded event rate to represent the inference signals.

To achieve this, we wire the network so as to keep gradient and inference signals separated. Specifically, all events are sent to somata up the hierarchy (Fig. 1A, orange links) while only bursts are sent down the hierarchy and these axons only target dendrites (Fig. 1A, yellow links), thus multiplexing feedforward signals with feedback signals. To ensure that the feedback signals are carrying credit information, we ensure the dendrites at the top of the hierarchy receive the classification error (Fig. 1B). The mathematical formulation of burst generation and plasticity are then chosen so as to approximate the backrpopagation of error algorithm for rate-based representations (see Supplementary Information of Ref. [10]). We note that the dendrites could in principle receive other signals than a supervised error signal. For instance, prediction errors signals in the dendrites would allow for the implementation of self-supervised algorithms [68]. We also note that the role of the precise structure of feedback connections has long been recognized as important and will be studied in Sect. 4.2 (Fig. 1C).

**Figure 1.**
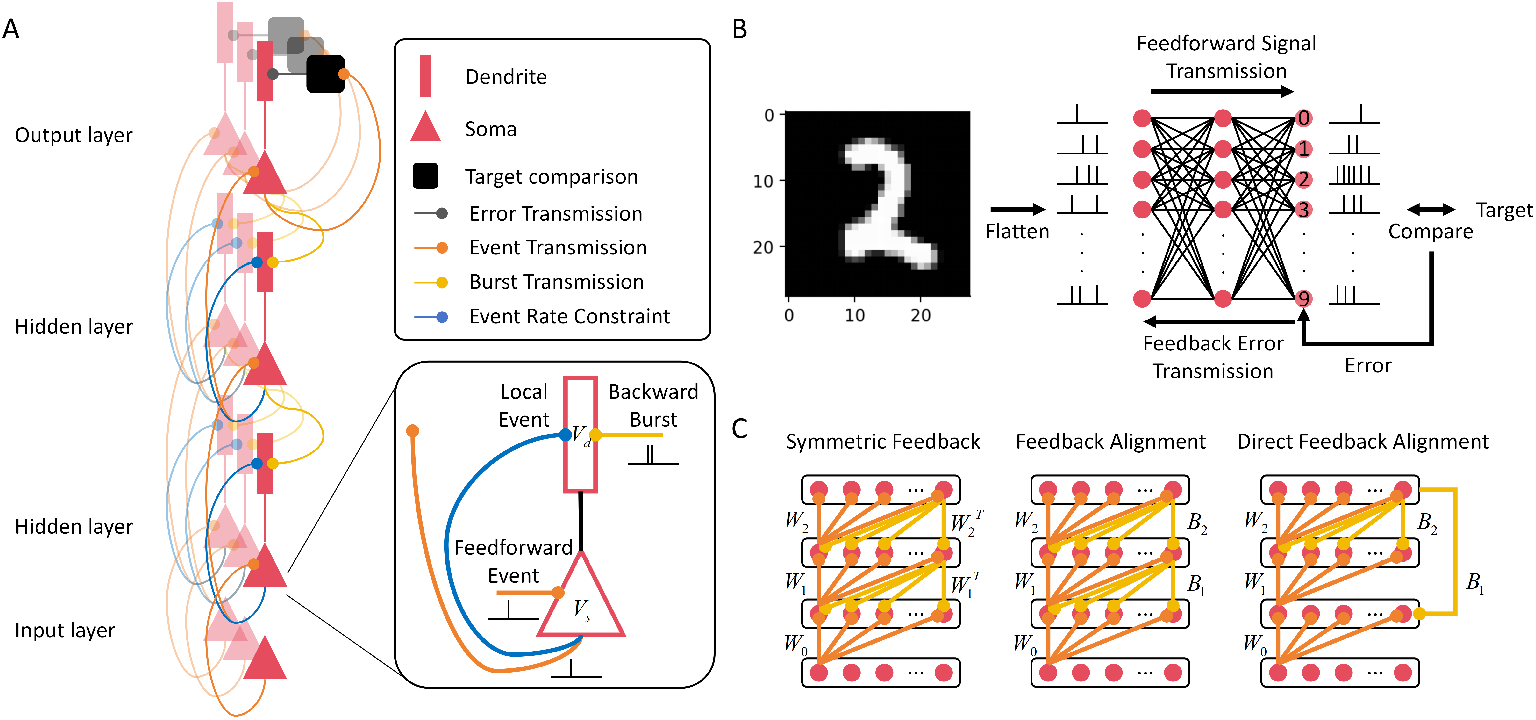
Overview of Burstprop connectivity and signal transmission: **A** Schematic of the core spiking neural network architecture. Each neuron is modeled as a two-compartment model with a dendrite (rectangle) and soma (triangle). The inputs to the dendrites modulate burst generation while the input to the soma modulate event generation, which can be either a single spike or a burst. Axons for each principal cell (orange and yellow) can connect locally, to the layer above, or to the layer below. Axons can selectively propagate either events (orange) or bursts (yellow). When connections are made to a lower-level unit, they target the dendritic compartment. When connections are made to a higher-level unit, they connect to a somatic compartment. Somatic compartment controls the rate of events being generated while the dendritic compartment controls the ratio of such events that are bursts. **B** Schematic of network for performing MNIST-like tasks. Images are flattened to a vector and pixel intensities are converted to spike train through Poisson firing. These form the input layer of the network whose detailed architecture is shown in A. A supervision is introduced at the readout layer where the firing in the desired unit is enforced by the creation of an error signal, injected in the readout units’ dendrites. **C** Schematic of the different types of feedback connectivity considered in this work. We used *W*_0_, *W*_1_ and *W*_2_ to label the feed forward weights from input, first layer or second layer, in the same order. These weights matrices are re-used for the feedback weights in the symmetric feedback scenario. In feedback alignement and direct feedback alignment, different weight matrices are used: *B*_1_, *B*_2_.

While Burstprop uses increases and decreases of burst probability to communicate positive and negative signs of the error signals, here we also considered an alternative model where the different sign of the error signal triggers different types of bursts. Physiologically, this ‘Signed Burst’ model could correspond to bursts of different durations engaging different short-term plasticity [35] and long-term plasticity. By construction, the signed burst model is more stable and less subject to drifts in plasticity than the original Burstprop model.

In Figure 2, we illustrate the two different types of bursting in with the imaginary case of a sinusoidal error signal. Errors are practically never sinusoidal but they can switch from positive to negative in the course of inference and learning samples. Whereas, in the unsigned burst model, the burstprobability oscillates around a fixed mean and it is by crossing this mean level that the neurons are able to represent positive and negative errors (Fig. 2B, left), in the signed burst model we have two burst probabilities, one for each type of burst, and the mean level is inconsequential for telling the sign of the error.

**Figure 2.**
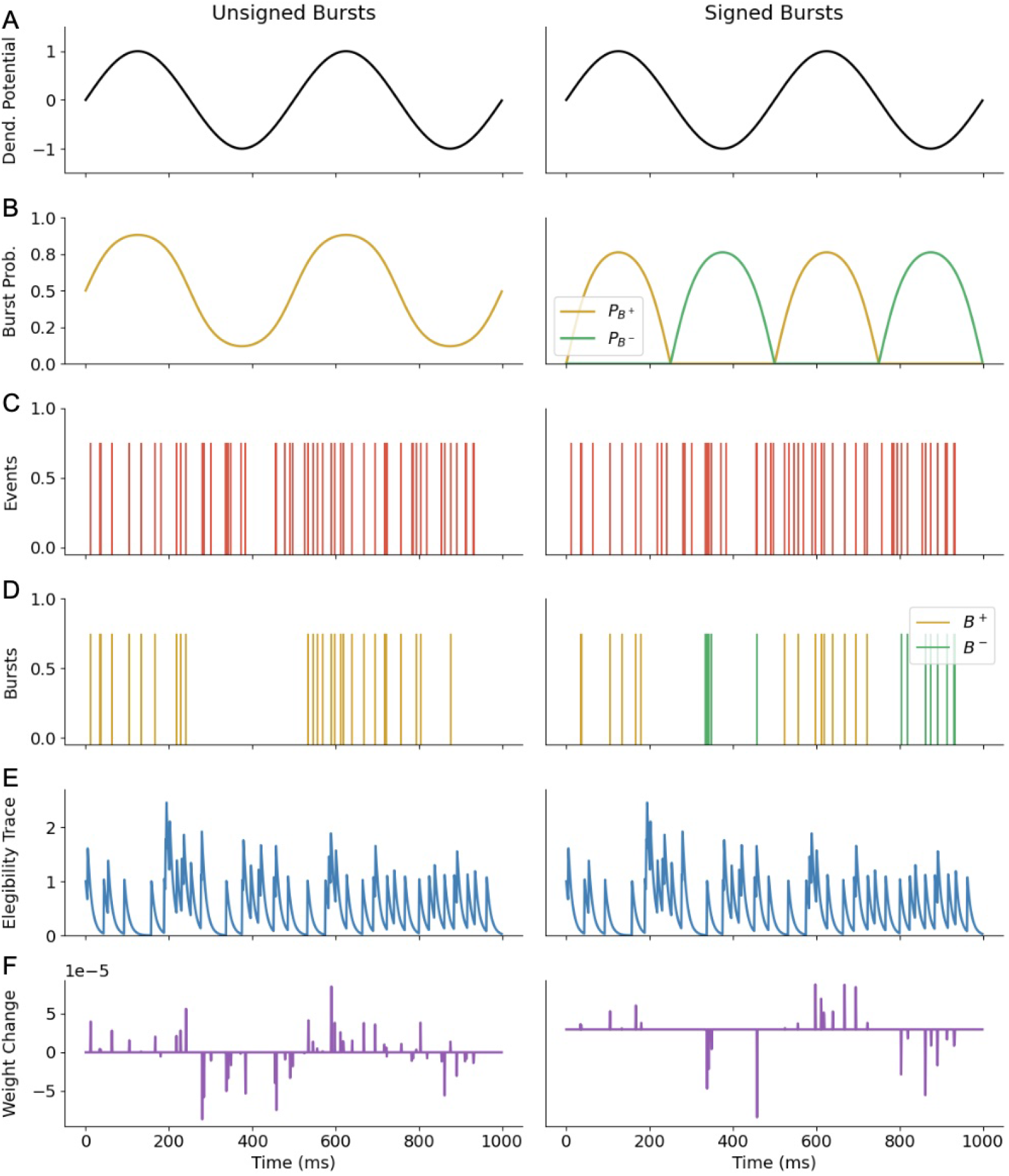
Dendritic potential steers direction of plasticity for synapses to neurons. A. Time-varying dendritic potential following a sine curve. B. Burst probabilities. Unsigned Burstprop, *P*_*B*_ = *σ*(*V*_*d*_). Signed Burstprop, *P*_*B*_+ = max(0, tanh(*V*_*d*_)) and *P*_*B*_ − = max(0, − tanh(*V*_*d*_)). C. Poisson spike train, with 50 Hz firing rate. D. Burst train generated from spike train and burst probability. E. Eligibility trace of a presynaptic neuron. F. Resultant synaptic plasticity.

### 4.1. Deep Learning on MNIST Dataset

To test the performance of our SNNs, we have trained both signed and unsigned versions of networks using symmetric feedback, varying width and depth on the rate encoded MNIST data (Fig. 3). We found by inspecting the resulting test-set errors that both signed and unsigned models were able to effectively train spiking neural networks with one layer with the signed model achieving 97.2 *±* 0.1% and 96.8 *±* 0.1% test classification accuracy respectively on a network with a single hidden layer of 100 neurons. For deeper networks, the signed model outperformed the unsigned one (Fig. 3A1), with the signed model achieving 95.9 *±* 0.1% accuracy compared to 64 *±* 1% accuracy for the unsigned model on a 4 hidden layer network. We did not observe an increase but rather a decrease in performance with the number of hidden layers. While there was no performance gain with depth, increased width was associated with an increase in performance (Fig. 3A2). Our algorithms could thus successfully train SNNs to perform MNIST with a performance that increases with layer width. Furthermore, we conclude that the signed burst model helps the algorithm to remain successful in deep networks.

**Figure 3.**
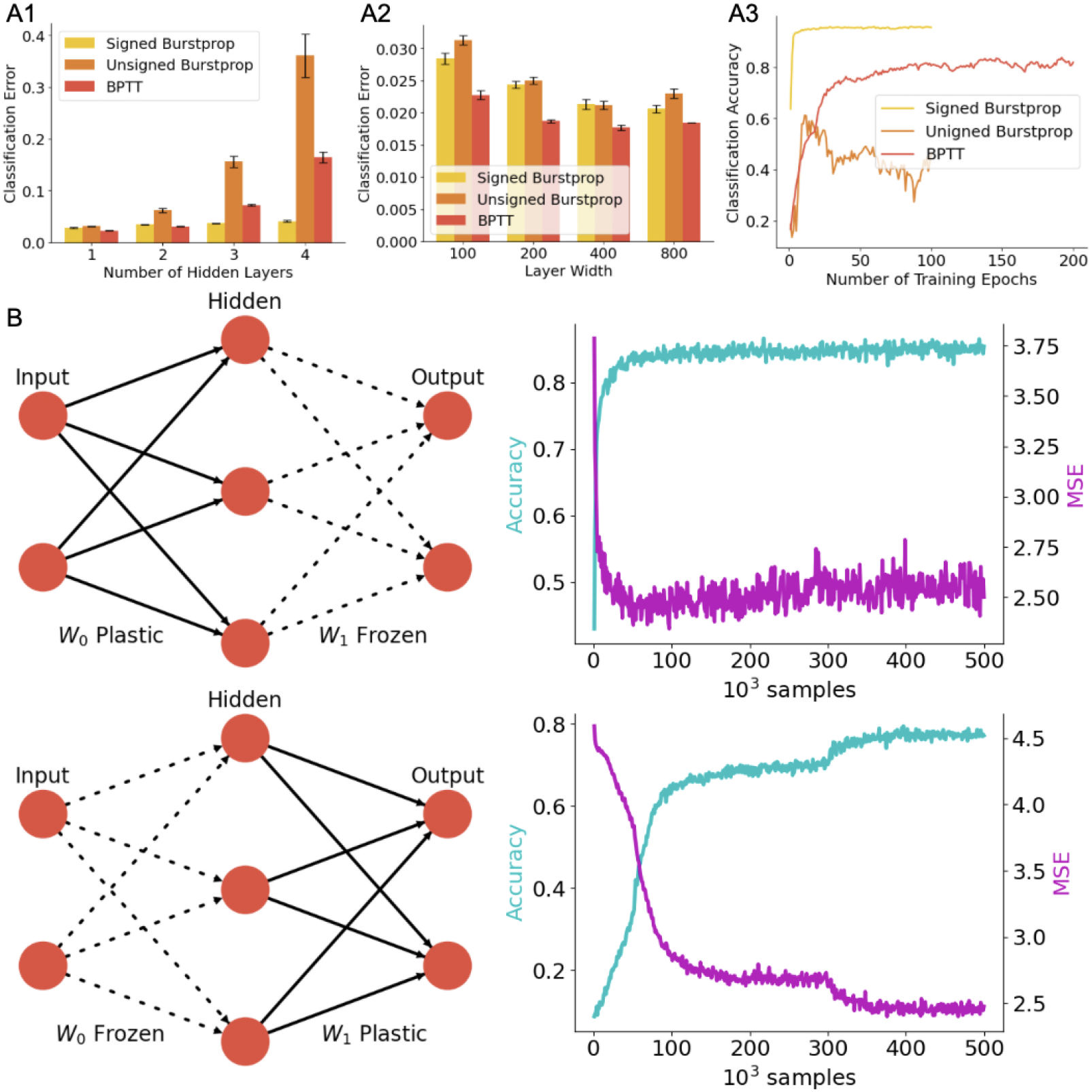
Burstprop coordinates multilayer learning to solve MNIST. A. Comparing test classification error on spiking MNIST dataset for Signed Burstprop, Unsigned Burstprop, and BPTT with symmetric feedback for networks with an increasing number of hidden layers, 100 neurons per layer (A1), and networks with increasing layer width, 1 hidden layer (A2). Error bars are standard error of the mean over 10 initializations. Burstprop algorithms were trained for 100 epochs and BPTT was trained for 200 epochs. A3. Example learning curves for each algorithm on a network of 4 hidden layers, each with 100 neurons. Error bars represent the standard deviation of minimum test error for 10 simulations. B. Signed Burstprop test accuracy and mean square error (MSE) per 10^3^ samples of a single hidden layer neural network of 100 hidden neurons. Row 1: *W*_0_ learns and *W*_1_ is frozen. Row 2: *W*_1_ learns and *W*_0_ is frozen.

To contextualize the learning ability, we compared the test-set prediction accuracy of our local learning algorithm to a state-of-the-art, non-local, BPTT-based learning method [21]. While burstprop aims to approximate the back-propagation of error algorithm, we could not directly compare with this algorithm as no good equivalent for SNN is commonly used. We found that in shallow networks, BPTT training applied on the same SNN architecture provided slightly better performance, with 97.7 *±* 0.1% accuracy on a network with a single hidden layer of 100 neurons. This improvement was maintained across the different network widths (Fig. 3A2). Networks trained with BPTT did not show a decrease in performance with network depth (84*±*1% accuracy on a 4 hidden layer network), an observation that echoes previous tests [66]. Because the choice of hyperparameters is likely to influence the comparison of these three algorithms, we conservatively conclude that SNNs with local burst-dependent plasticity rules can achieve good performance on MNIST, even with a small number of neurons and with multiple layers.

That the network performance does not increase with network depth may suggest that the algorithm is only training the output weights. To investigate this, and to determine whether the algorithm is effectively assigning credit to earlier layers, we examine how freezing either the output weights or the input weights affects learning in a single hidden layer network. If learning were limited to the output layer, freezing plasticity in this layer should result in the network being unable to learn. Instead, what we see is that the network remains able to learn effectively when the output layer is frozen (Fig. 3B). To take another approach, forcing the plasticity to only occur in the output weights should preserve the performance if credit signals were not reaching the lower-order connection. We found that freezing the output weights has a smaller effect on learning capabilities than freezing the input weights. Freezing the input weights made learning much slower (Fig. 3). Together, our burst-dependent SNN can successfully propagate credit-carrying information to neurons and accurately steer plasticity beyond the output layer. This suggests that our algorithm is effectively approximating backpropagation in the way it assigns errors.

### 4.2. Feedback Alignment

While we have shown that Burstprop can successfully train an SNN on MNIST, we did this by enforcing symmetry between feedforward and feedback weights. To allow for neuromorphic learning, and to address biological plausibility, our algorithm must remain effective with weight-transport-free learning. We thus tested the performance of Burstprop on our spiking MNIST dataset using feedback alignment (FA) and direct feedback alignment (DFA), two algorithms that avoid the weight-transport problem by using random but fixed feedback weights (see Methods and Figure 1C). If Burstprop can learn from the data while using random but static feedback weights, this would suggest that weight transport is not necessary for learning in SNNs. We found that both signed (Figure 4B1) and unsigned (Figure 4B2) versions of the algorithm can achieve good performance, achieving 95.6 *±* 0.1% accuracy and 85 *±* 1% accuracy respectively on a 4 hidden layer network with feedback alignment. The decrease in performance with the number of layers was lessened by switching to FA for the unsigned burst model but unaltered for the signed burst model.

**Figure 4.**
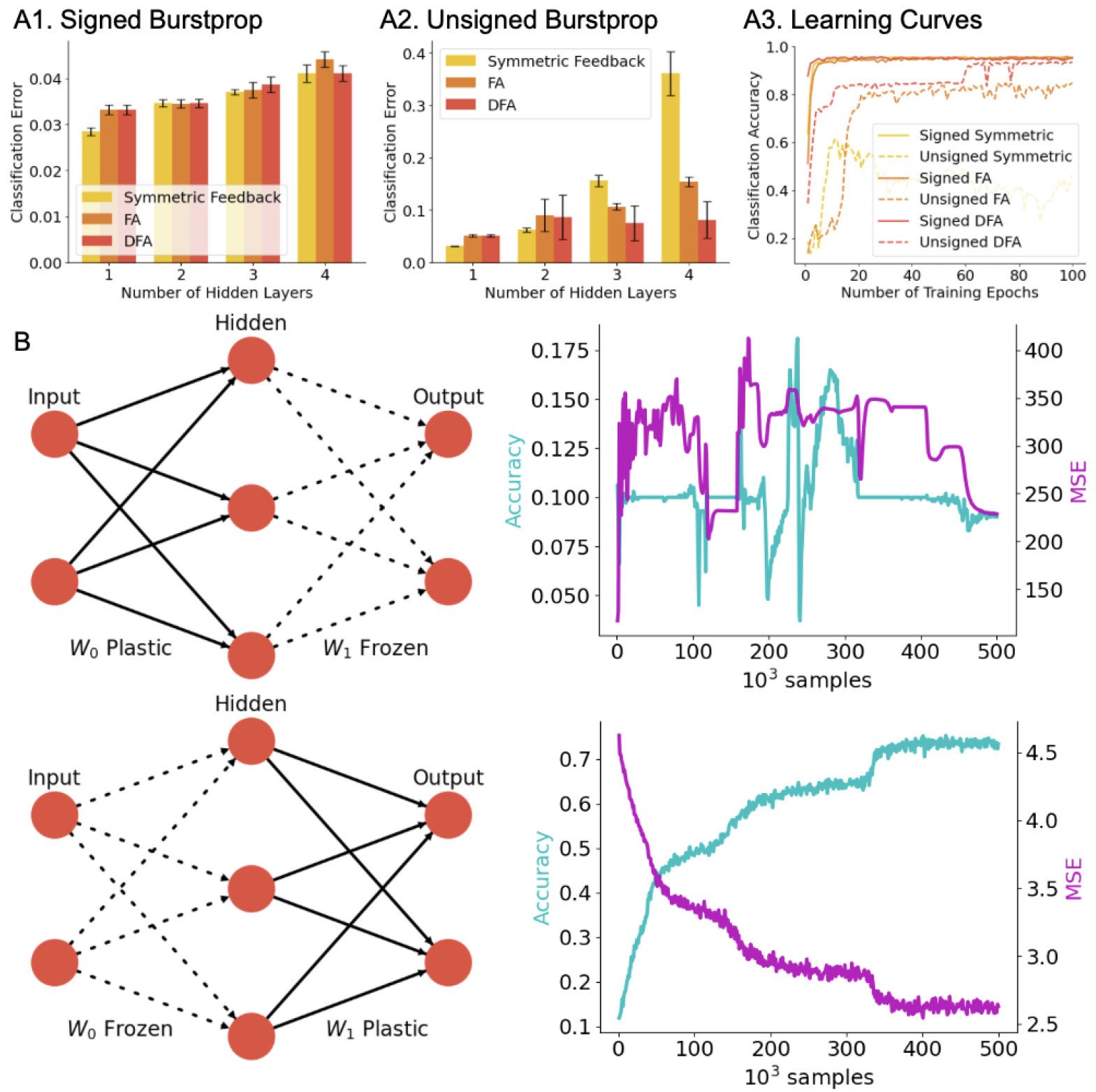
Burstprop learns with feedback alignment methods: A. Comparing test classification error on spiking MNIST dataset for different feedback error transportation methods with the Signed Burstprop (A1) and Unsigned Burstprop (A2) algorithms with increasing number of hidden neurons, 100 neurons per layer. Each trained for 100 epochs. Error bars are standard error of the mean over 10 initializations. A3. Example learning curves for each algorithm on a network of 4 hidden layers, each with 100 neurons. B. Signed Burstprop with FA test accuracy and mean square error (MSE) per 10^3^ samples of a single hidden layer neural network of 100 hidden neurons. Row 1: *W*_0_ learns and *W*_1_ is frozen. Row 2: *W*_1_ learns and *W*_0_ is frozen.

Next, we again investigated how freezing either the first or second layer impacts learning. In principle, learning the set of weights should not be possible if the higher-order layers have not had a chance to learn and align with the feedback [39]. Accordingly, if we freeze the output weights, learning should be strongly affected. This is indeed what we see (Fig. 4B), whereby freezing the output weights in an FA scenario completely blocked learning. Thus using random but fixed feedback weights can implement weight-transport-free learning in SNNs using local learning rules.

### 4.3. Testing EMNIST and FMNIST

MNIST is recognized as an exceedingly easy task for in machine learning as a linear classifier, especially paired with a sufficiently expressive input basis, already achieves over 90% accuracy [67]. In addition to the weight-freezing manipulation, we also tested the learning algorithm on other, more difficult but closely related tasks. As our algorithm is meant to approximate backpropagation, we do not expect that temporal tasks would be a good choice as these tasks typically require the use of biologically challenging algorithms such as backprop-through time [21]. Because of the computational load of simulating SNNs on traditional hardware, we restricted our tests to EMNIST and FMNIST. Averaging over 5 different initialization with different initial degree of connection probability, we found that while MNIST achieved an accuracy of 91.0 *±* 2.2 %, the network achieved 91.1 *±* 1.5 % on EMNIST and 82.6 *±* 0.5 % on FMNIST.

We found that the performance was relatively better for EMNIST [69] and FMNIST [70]. The accuracy reached for EMNIST was 91.8 % and 81.2 % for FMNIST, that is higher or equal accuracies than for MNIST. This is to be compared with linear classifiers which achieve over 90 % on MNIST [67] and 56 % on EMNIST [69] and 83 % on FMNIST [70] The performance of traditional neural networks also tends to be lower on EMNIST and FMNIST than on MNIST [71, 72].

### 4.4. Sparse connectivity

Because many hyper-parameters can have a profound influence on performance, we wanted to test our algorithm further. We chose to focus on the role of the sparsity of the initial connectivity because 1) sparsity can reduce catastrophic forgetting [73] and 2) cortical neural networks are exceedingly sparse [74]. We simulated learning on all three benchmark datasets while varying the sparsity of the initial connectivity from 0 to 0.8 (Table 1). We found that sparser initializations of the networks had a non significant tendency to degrade the test accuracy with increasing sparsity (Pearson *r* =-0.43, *P* = 0.11).

**Table 1.** Test performance under different conditions of sparsity *p*–single simulation per condition. A high value of *p* means the network was initialized with many weights put to zero.

|  | MNIST |  | EMNIST |  | FMNIST |  |
| --- | --- | --- | --- | --- | --- | --- |
|  | accuracy | error | accuracy | error | accuracy | error |
| $p = 0.0$ | 0.945 | 1.771 | 0.941 | 1.803 | 0.836 | 2.193 |
| $p = 0.2$ | 0.941 | 1.877 | 0.933 | 1.858 | 0.838 | 2.255 |
| $p = 0.4$ | 0.929 | 1.912 | 0.917 | 1.773 | 0.83 | 2.332 |
| $p = 0.6$ | 0.926 | 1.758 | 0.847 | 2.119 | 0.813 | 2.335 |
| $p = 0.8$ | 0.811 | 2.192 | 0.918 | 1.923 | 0.812 | 2.179 |

### 4.5. Low-Resolution Synaptic Weights

An important consideration for implementing SNNs in neuromorphic hardware is the number of distinct values that synaptic weights can take [75]. Spiking neuromorphic hardware may only be able to support small bit resolutions for their weights, in the range of 4 to 8 bits [75]. Here we investigate the impact of decreased synaptic weights on learning performance. Figure 5 shows that learning performance decreases with weight resolution. Still, even with only 4-bit resolution, learning is achieved with over 90% classification accuracy. This result indicates that Burstprop is robust to low bit resolution in the synaptic weights.

**Figure 5.**
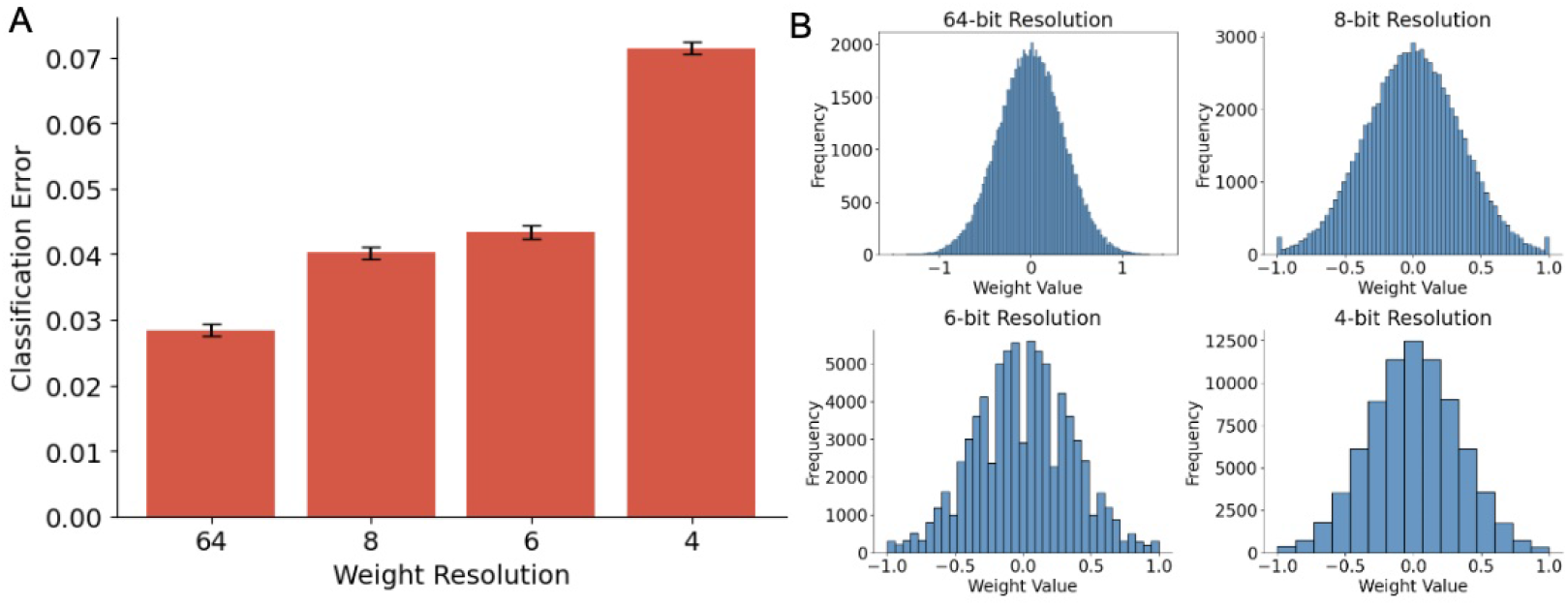
Burstprop is robust to low-resolution synaptic weights. A. Comparing test classification error on spiking MNIST dataset for networks of a single hidden layer of 100 neurons, trained with Signed Burstprop, with varying weight resolutions. Error bars are standard error of the mean over 10 initializations. B. Initial weight distributions of input weights for varying weight resolutions.

## 5. Discussion

We have implemented a spiking version of the Burstprop algorithm that uses local learning rules and marked point processes for differential communication to different post-synaptic targets. Our algorithm improved on the previous efforts solving XOR with over a few thousand neurons to solving more complex tasks with a few hundred neurons. We found that our SNN using a local, burst-dependent learning rule could achieve a performance similar to that of BPTT. By experimenting with freezing weight layers during learning, we conclude that the algorithm can accurately perform credit assignment beyond the output layer. Training the algorithm with FA and DFA also showed that Burstprop can successfully perform weight-transport-free learning. The results further show the algorithm is robust to low-bit resolution synaptic weights. These results indicate that burst-dependent communication and plasticity can form an effective, biologically inspired, local learning method for SNNs that may be useful for neuromorphic learning.

We have shown that feedback alignment methods can be used effectively with SNNs using Burstprop to solve weight transport between feedforward and feedback weights. Our signed burst algorithm, however, requires some symmetry between two sets of feedback weights. Biological neurons would need two different synapses to communicate different firing patterns. These symmetric pathways are required to transmit positive and negative error signals. In our experiments, we have been enforcing this symmetry through weight transport, however, many methods could be implemented to align these weights. Further work must be done to find a suitable solution to address this problem. This problem is addressed in previous work [11] by adding a learning rule to feedback synapses, ensuring they converge.

Another weakness of our algorithm is that its fitting capabilities were not improved by increasing network depth. This is common for spiking neural networks [66, 50] and it is unclear whether this is related to the input encoding or could be a fundamental limitation of the algorithm. While neural networks often rely on deep structures to fit complex functions, even a single-layer spiking neural network is a universal function approximator so width could be used to compensate for depth in some applications.

An important limitation of our results worth mentioning is that we limit our experimentation to MNIST-like datasets. MNIST itself is a fairly simple benchmark problem for supervised learning algorithms and many non-neural network algorithms reach high test accuracy on this task. For instance, a linear classifier can achieve 7.6% error [67], a random forest algorithm can achieve 2.4% error, and an SVM can achieve 0.56% error [76]. The goal of this work was not to achieve state-of-the-art performance on this benchmark, but to show that Burstprop can be applied to solve a more complex learning task than XOR. Further experimentation with more complex problems is required to see how that algorithm scales.

Current neuromorphic hardware typically emulates leaky integrate-and-fire-like neurons, and rarely implements dendritic compartments [77, 78], bursting or burst-dependent plasticity. Our work suggests that developping such a device can allow spiking neural networks to learn on chip, which should improve substantially both the energy reauirements and the speed of training.

